# Secure Wavelet Matrix: Alphabet-Friendly Privacy-Preserving String Search

**DOI:** 10.1101/085647

**Authors:** Hiroki Sudo, Masanobu Jimbo, Koji Nuida, Kana Shimizu

## Abstract

**Motivation:** Privacy-preserving substring matching is an important task for sensitive biological/biomedical sequence database searches. It enables a user to obtain only a substring match while his/her query is concealed to a server. The previous approach for this task is based on a linear-time algorithm in terms of alphabet size |Σ|. Therefore, a more efficient method is needed to deal with strings with large alphabet size such as a protein sequence, time-series data, and a clinical document.

**Results:** We present a novel algorithm that can search a string in logarithmic time of |Σ|. In our algorithm, named secure wavelet matrix (sWM), we use an additively homomorphic encryption to build an efficient data structure called a wavelet matrix. In an experiment using a simulated string of length 10,000 whose alphabet size ranges from 4 to 1024, the run time of the sWM was an order of magnitude faster than that of the previous method. We also tested the sWM on all sequences of one protein family in Pfam (9,826 residues in total) and clinical texts written in a natural language (77,712 letters in total). By using a laptop computer for the user and a desktop PC for the server, we found that its run time was ≈ 2.5 s (user) and ≈ 6.7 s (server) for the protein sequences and ≈ 10 s (user) and ≈ 60 s (server) for the clinical texts.

**Availability:** https://github.com/cBioLab/sWM

## 1 Introduction

Privacy-protection is an emerging problem in the field of life science where many data are private, such as clinical documents, health records, personal genomes and protein sequences. Currently, the most common approach for protecting such data is to limit access by segregating the data, which eventually deteriorates their value. To overcome this problem, a technology is eagerly demanded that enables such sensitive data to be searched while maintaining individual privacy.

In this study, we consider a typical scenario where the user wants to search on a text database but does not wish to show his/her query to the server, and the server wants to return only the search result and does not want the user to obtain any other information. In a previous study aiming to achieve such a scenario, Freedman *et al.* (2005) introduced a keyword search, in which both the server and the user have a set of keywords and only the user knows common keywords. Bruekers *et al.* (2008) also developed a similar approach and applied it to a genetic test. There is another line of studies aiming to evaluate similarity of a query and each database entry by computing a Jaccard Index of two keyword lists. Applications have been developed for a DNA sequence search (Baldi *et al.*, 2011; Cristofaro *et al.*, 2013) and a chemical compound search (Shimizu *et al.*, 2015). In addition to those keyword-based searches, a privacy-preserving substring search is also an important problem in bioinformatics. In a substring search, the server has a long text and the user has a query (a relatively shorter text), and only the user knows the positions where the query matches to the server’s text. In previous work (Katz and Malka, 2010), a DNA sequence search was demonstrated using a relatively time and communication-size consuming approach (Yao, 1986) that mainly targets an exact match. Recent studies by Vergnaud (2011); Baron *et al.* (2013) presented more efficient methods to find matches that have a small fixed Hamming distance. While those studies aimed to evaluate similarity between an entire query and a server’s text, another study (Shimizu *et al.*, 2016) aimed for a variable-length prefix/suffix match either on a regular text or a set of aligned texts such as SNP sequences (e.g., it enables the user to find only occurrences of the longest prefix/suffix match), which is also useful for various practical settings. In principle, the method in this previous study (called PBWT-sec) can be applied for any type of text data. However, it does not perform sufficiently for practical problems when it is used for searching on a text with a large alphabet size |Σ|, such as a protein sequence (|Σ| = 21) and clinical documents (e.g., |>Σ| = 36 for Roman alphabet and Arabic numerals, |Σ| = 83 for Japanese alphabet), because the algorithm was originally designed for haploid genome sequences (|Σ| = 2) and its time complexity is linear to |Σ|.

To overcome this critical drawback, here we present an efficient algorithm named the secure Wavelet Matrix (sWM) that builds an efficient data structure called a wavelet matrix (Claude and Navarro, 2012) by using additively homomorphic encryption. As we will later discuss in Section 3.5 and Section 4, the sWM is an order of magnitude faster than the PBWT-sec in terms of alphabet size, without losing any advantages of the previous framework used in PBWT-sec, and thus it can practically deal with various types of text data. We will also describe a detailed algorithm to use sWM for a full-text search based on FM-Index (Ferragina and Manzini, 2000).

The rest of the paper is organized as follows. In Section 2, we describe the main idea of our approach, which is followed by Section 3 where the detailed algorithms of sWM and its application to full-text search are provided. We also describe the complexity of the algorithms and compare it with that of previous methods. In Section 4, we evaluate the performance of our method both on a simulated dataset and two real datasets (protein sequences and clinical texts) and compare our method with a previous approach and a baseline approach. Finally, we conclude our study in Section 5.

## 2 Approach

### 2.1 Problem setting

In this study, we assume a case in which a user has a query text ***q*** and a database holder has a long text *S*, and the user wants to know only the longest prefix of ***q*** that matches to *S*. As explained in Section S1 in the supplementary material, the proposed method is easily modified to find the longest prefix that matches more than *ϵ* different positions in *S* given the threshold *ϵ*, which may fit more practical applications. For ease of explanation, we describe a series of algorithms specialized for searching on a single long text, but the same approach is able to search on other types of text data such as a set of aligned texts by modifying it slightly.

### 2.2 Overview

To explain the central idea of our approach, we will first briefly describe PBWT-sec (Shimizu *et al.*, 2016). PBWT-sec is designed to search on an iteratively queriable data structure such as Burrows-Wheeler Transform (BWT), in which a database text is efficiently indexed. During the search using such a data structure, a match between a query and a database text is reported as a left-close and right-open interval [*f, g*) on the auxiliary data, usually a sorted order of the original text, as shown in Figure 1. We do not provide a detailed mechanism of the data structure. However, to understand the key idea, two points are sufficient to know. (1) The lower bound *f* and upper bound *g* of the interval are computed by a function called RankCF. (2) An interval [*f*_*k*+1_, *g*_*k*+1_) for a prefix match of length *k* + 1 between a query text ***q*** and a database text *S* is computed in the form of:

**Figure 1:**
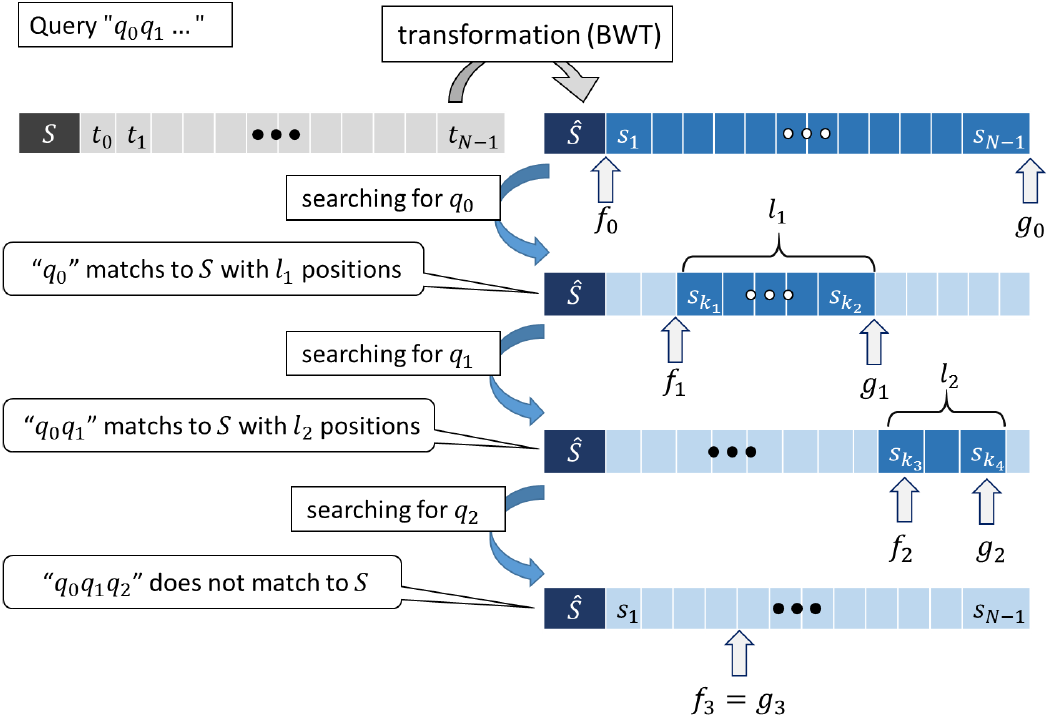
Schematic view of search on recursive data structure such as BWT. Each match is represented by interval [*f,g*), and interval is updated by previously computed interval.

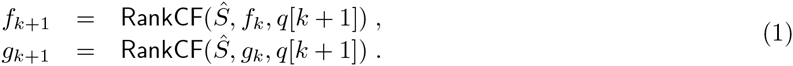

where *Ŝ* is an auxiliary text such as BWT of *S*, and ***q***[*k* + 1] is the (*k* + 1)-th letter of ***q***.

As we explain in detail later in this section, `RankCF` is pre-computable and stored in a long lookup table ***υ***, and thus [*f, g*) can be obtained by referring to υ:

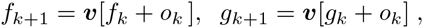

where *o_k_* returns proper offsets necessary for storing all pre-computed values. Therefore, the interval for a prefix match of length *k* + 1 can be obtained by referring to the same table ***υ*** recursively as follows.

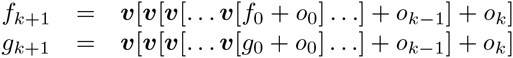

To enable the user to refer the lookup table held by the server, a cryptographic technique called a *Recursive Oblivious Transfer* (ROT) protocol is used, by which ***υ***[***υ***[… ***υ***[*i*]…]] is returned to the user when the user inputs an index *i* and the server inputs a vector ***υ*** to the protocol. In the process of the ROT protocol, the user and the server repeat communication, and in each round, the user obtains (***υ***[*x*] + *r*)_mod *N*_ where *x* is the user’s input, *r* is the server’s input (a random factor) and *N* is the length of ***υ***. All the user’s inputs are encrypted by the user’s secret key before it is sent to the server, and the server computes return values on the basis of the user’s encrypted values without decryption. Since the server does not have the secret key, the user’s inputs are not leaked to the server. On the other hand, the user decrypts the returned values, but cannot see ***υ***[*x*] because it is randomized by the server’s private value *r*. The key feature of ROT is that when the user inputs previously returned value (***υ***[*x*] + *r*)_mod *N*_, the server properly removes *r* from the user’s input and returns (***υ***[***υ***[*x*]] + *r^′^*)mod *N* where *r^′^* is another random factor. At the end of the protocol, the user can obtain a non-randomized result if the server does not input a random factor. In this way, `ROT` enables the user to refer to ***υ*** without leaking any intermediate information. (See Section 3.2 for more details about the algorithm of `ROT`.) By using this property and an additional cryptographic technique, the PBWT-sec can safely compute *f* and *g* until it finds the longest prefix (i.e., *g − f* becomes 0). Figure 2 illustrates how the user and the server communicate when computing the interval.

**Figure 2:**
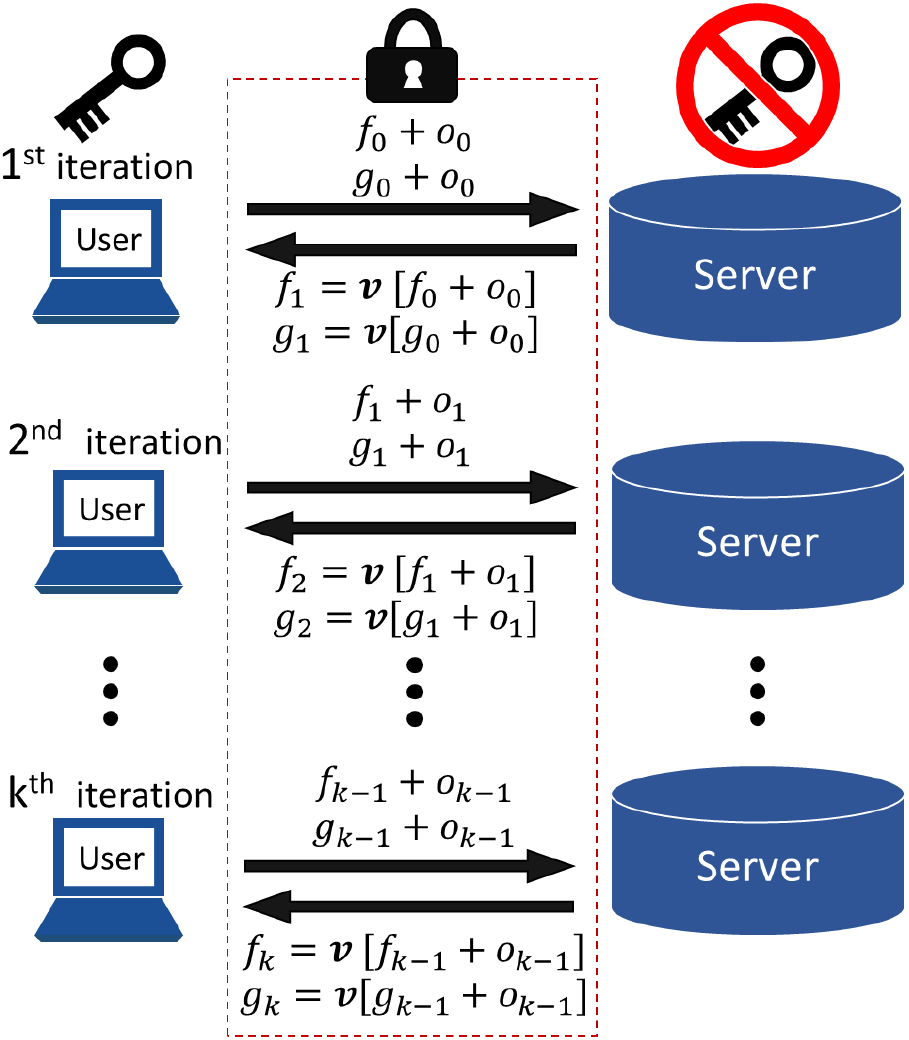
Schematic view of communication between user and server for computation of interval [*f, g*). Lower bound *f_k_* and upper bound *gk* are obtained by *f*_k+1_ = ***υ***[*f_k_*+ *o_k_*] and *g_k+1_* = ***υ***[*gk* + *o_k_*], respectively. All intermediates *f_0_*,…, *f_k_* and *g_0_*,…, *g_k_* are concealed by using ROT protocol.

The `ROT` task is a computational bottleneck of the PBWT-sec, and the time complexity is linear to the length of ***υ***. The goal of this study is to improve the time complexity of the `ROT` task. Let us give more details about ***υ***and `RankCF`, which are key building blocks of `ROT`. Given an index *p*, a character *c* and a sorted text *Ŝ* (such as **BWT**) of length *N*, `RankCF`(*Ŝ*, *p*, *c*) is defined as follows.

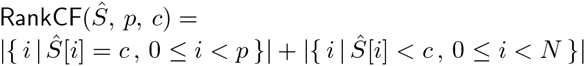

We denote *Ŝ*[*i*] < *c*, when a letter *Ŝ*[*i*] is lexicographically smaller than a letter *c*. For example, if *Ŝ* =“MDGGIPQAGG”, *p* = 5 and *c* =‘G’, `RankCF`(*Ŝ*,5,‘G’) becomes 4 because the two leftmost ‘G’s are within the first five letters, and ‘D’ and ‘A’ are lexicographically smaller than ‘G’.

The straightforward method to store all outputs of `RankCF` for the *Ŝ* of length *N*, 0 ≤ *p* ≤ *N*, and *c* ∈ Σ is to create a lookup table of length *N* × |Σ|. In fact, **PBWT**-sec designs ***υ*** in this simple way, and thus its computational complexity is linear to |Σ|.

To reduce the total cost for the `ROT` task, we propose a novel approach using *wavelet matrix* (Claude and Navarro, 2012) to design ***υ*** very efficiently.

### 2.3 Wavelet Matrix

A wavelet matrix (WM) is an efficient data structure that supports a wide range of query operations such as `RankCF`. It achieves logarithmic-time complexity in terms of alphabet size |Σ| while keeping space complexity close to information-theoretic lower bound. Given a text *T*_0_, the key feature of WM is to encode each letter in a binary form and to compute `RankCF` bit by bit from the least to the most significant bit in order to compute final RankCF for *T*_0_. Let us describe this process in more detail. A WM algorithm creates a bit array *B*_0_ = *b*^0^(*T*_0_[0]),…, *b*^0^(*T*_0_[*N* − 1]), where *b^i^*(*c*) denotes *i*-th bit of the binary encoding of *c*. The algorithm obtains new text *T*_1_ by sorting *T*_0_ in accordance with *B*_0_, and another bit array *B*_1_ = *b*^1^(*T*_1_[0]),…, *b*^1^(*T*_1_[*N* − 1]) is created in the next step. *B_i_* is created from *B_i−1_* in a similar manner until *i* reaches *λ* − 1 when a bit length *λ* = [log |Σ|]. There is the following recursion relationship:

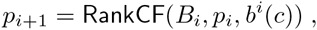

which leads to

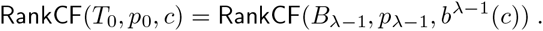

Figure 3 illustrates an example of a search on WM.

**Figure 3:**
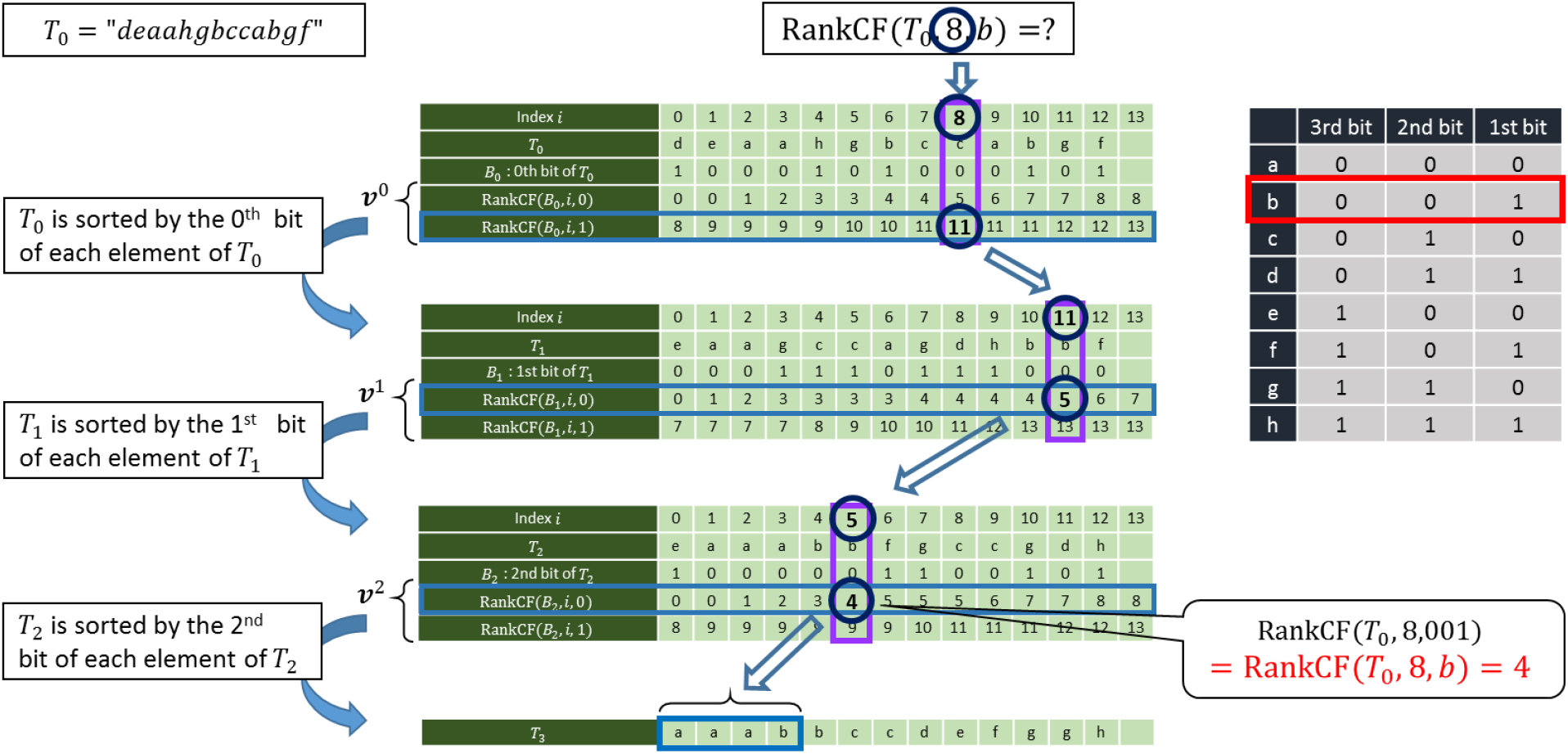
Example of search on wavelet matrix. `RankCF`(*T*_0_, 8, *b*) is obtained by `RankCF`(*B*_0_, 8, *b*^0^(*b*)), which returns 11; `RankCF`(*B*_1_, 11, *b*^1^(*b*)), which returns 5; and `RankCF`(*B*_2_, 5, *b*^2^(*b*)), which returns 4.

### 2.4 Efficient design principle of lookup table *υ*

Here, we explain how to design the lookup table ***υ*** efficiently. As described in Section 2.3, `RankCF`(*T*_0_, *p*_0_, *c*) is computed by repeating `RankCF` on auxiliary bit-arrays *B*_0_,…, B_*λ*−1_. In our approach, we create a set of sub-lookup tables ***υ^i^*** for *i* = 0,…, *λ* − 1, each of which corresponds to a bit-array *B_i_* as follows, and use `ROT` to refer to ***υ^i^***.

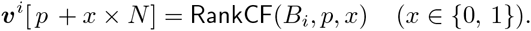

The outline of our method sWM is as follows. Note that the goal of sWM is to compute `RankCF`(*T_0_*, *p_0_*, *c*).

**Step 1** The server creates ***υ***^0^,…, ***υ^λ−1^*** from ***B_0_***,…, ***B_λ−1_***.

**Step 2** The user’s initial input to `ROT` is an encrypted *p_0_* + *o_0_*. The offset is *o_0_* = 0 when 0-th bit of user’s character *c* is 0, otherwise *o_0_* = *N*.

**Step 3** The server’s initial inputs to `ROT` are an encrypted ***υ***^0^ and a random factor *r*_0_.

**Step 4** The user obtains 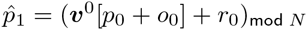 from `ROT`.

**Step 5 for** *i* = 1,…, *λ* − 1

- The user inputs encrypted 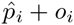 to `ROT`.
- The server inputs *r*_*i*−1_ and a new random factor *r_i_* and ***υ^i^*** to compute 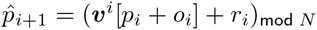 in an encrypted form.
- The user obtains 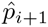.

By the above protocol, only the user can safely obtains (`RankCF`(*T*_0_, *p*_0_, *c*) + *r_λ−1_*)_mod *N*_. Note that the user can obtain `RankCF`(*T_0_*, *p_0_*, *c*) if the server sets r_*λ*−1_ = 0. Since the length of each sub-lookup table is 2*N* and s**WM** repeats `ROT` for *λ* times, the total time complexity becomes *O*(*N* log |Σ|), which is an order of magnitude better than the previous approach’s time complexity *O*(*N*|Σ|). Figure 4 illustrates the design principle of ***υ*** for s**WM** and that for **PBWT**-sec. We will describe the sWM algorithm in more detail in Section 3.3.

**Figure 4:**
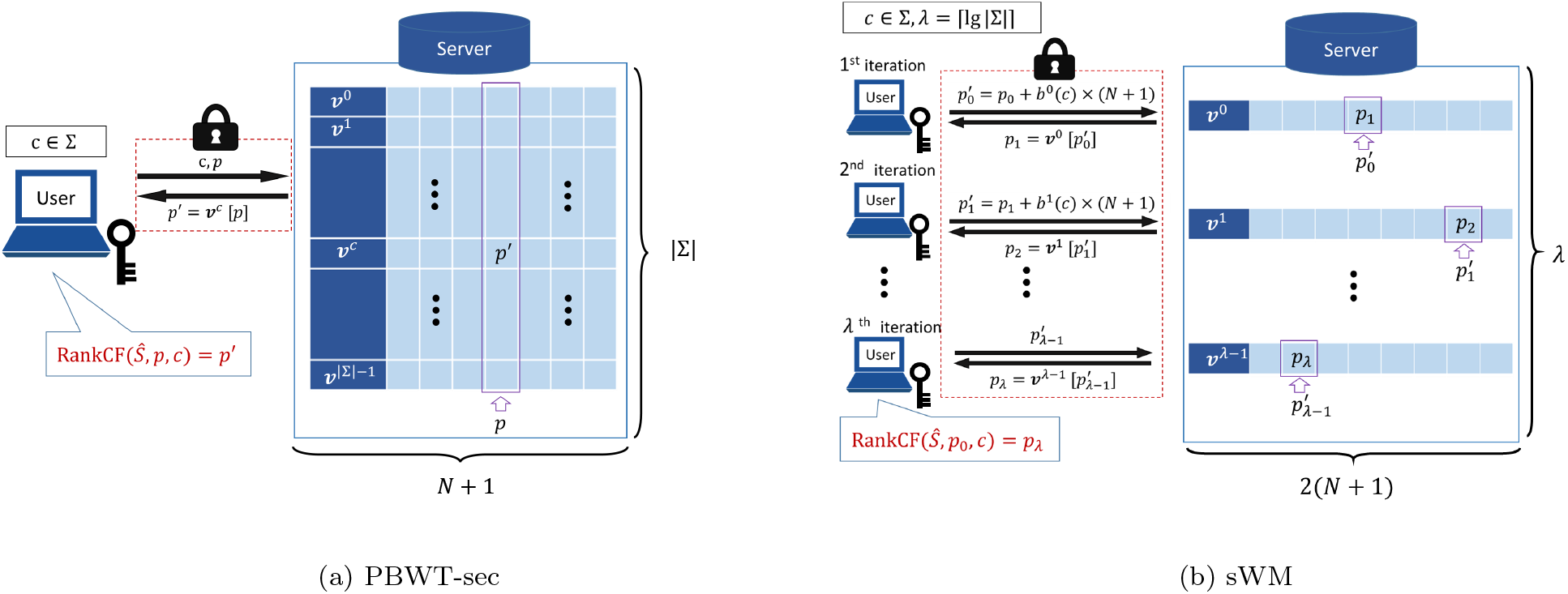
Schematic view of updating each bound of interval in (a) PBWT-sec (previous method) and (b) secure wavelet matrix (proposed method). In PBWT-sec, user sends *c, p* in such a way that only single communication occurs. Server returns ***υ***^*c*^[*p*] in encrypted form by scanning lookup table of length *N*|Σ|. Therefore, the time complexity of this task becomes *O*(*N*|Σ|). In sWM, log |Σ| communications occur to obtain the same information. For each communication, server returns 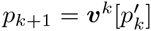 by scanning sub-lookup table of length *N*. Therefore, time complexity of total communications becomes *O*(*N* log |Σ|).

## 3 Method

As described in Section 2, the s**WM** is designed on the basis of the `ROT`, which we implemented by using additively homomorphic encryption. In this section, we describe those cryptographic building blocks and provide more details about the s**WM** algorithm.

### 3.1 Additively Homomorphic Encryption

Additively homomorphic encryption is a kind of public-key encryption scheme that enables addition of *encrypted* values to be computed. A public-key encryption scheme consists of three algorithms: the key generation algorithm **KeyGen** generates a public key **pk** and a secret key **sk**; the encryption algorithm Enc generates a ciphertext **Enc**(*m*) of message *m* under the given **pk**; and the decryption algorithm **Dec** computes the decryption result of a ciphertext under the given **sk**. An additively homomorphic encryption scheme also has the following additively homomorphic functionalities:

- An operation **Enc**(*m*_1_) ⊕ **Enc**(*m*_2_) to generate **Enc**(*m*_1_ + *m*_2_) from two given ciphertexts **Enc**(*m*_1_) and **Enc**(*m*_2_) of integer messages *m*_1_ and *m*_2_, without knowing *m*_1_, *m*_2_, or the secret key.
- An operation *e* ⊗ **Enc**(*m*) to generate **Enc**(*e • m*) from a given ciphertext **Enc**(*m*) and an integer *e*, without knowing *m* or the secret key (in particular, **Enc**(−*m*) can be computed by the operation).

We suppose that the scheme used in this study is semantically secure; that is, no information in the original message can be learned from a ciphertext (Goldwasser and Micali, 1984). Examples of other such schemes are the Paillier cryptosystem (Paillier, 1999) and the “lifted” version of the ElGamal cryptosystem (ElGamal, 1985), where the second operation ⊗ can be achieved by iteration of the first operation ⊕.

### 3.2 Recursive Oblivious Transfer

The `ROT` protocol consists of three sub-modules.

- `PrepQuery` 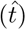 is a user-side sub-module. It takes an index 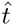 and returns an encrypted query. In fact, the query is in the form of a vector of ciphertexts for the further process of `ROT`. However, we do not go into detail about the specifications of `ROT` and we abuse 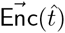 to denote an encrypted query.
- `RecQuery` 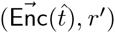 is a server-side sub-module. It takes a user’s input 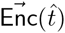and a random factor *r*^′^ that was used to compute 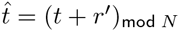 in the previous round and removes *r*_′_ to return 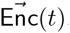.
- `RanOT` 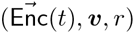 is a server-side sub-module. It is a computationally dominant part and computes an encrypted and randomized result **Enc**((***υ***[*t*] + *r*)_mod *N*)_ from 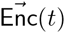, the server’s lookup table ***υ*** and a random factor *r*.

#### Algorithm 1

Detailed description of secure Wavelet matrix

- Public input: Problem size *N*; alphabet Σ; public key `pk`
- Private input of user: An index *p*_0_, a query character *c* ∈ Σ, private key `sk`
- Private input of server: A database text *T*_0_

1. (*Server initialization*)

a. Create auxiliary bit-arrays *B*_0_,…, *B*_*λ*−1_ from *T*_0_ **for** (*k* = 0,…, *λ* − 1) **do** 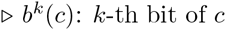

- *B_k_* = *b^k^* (*T_k_* [0]),…, *b^k^* (*T_k_* [*N* − 1])
- Sort *T_k_* by *b^k^* (*T_k_* [·]) to generate a sorted string *T*_*k*+1_ **end for**
b. Create sub-lookup tables ***υ***^0^, ***υ***^1^,…, ***υ***^*λ*−1^ ***υ^k^*** [*i* + *x* × *N*] = `RankCF`(*B_k_, i, x*) (0 ≤ *k* < *b*, 0 ≤ *i* ≤ *N, x* ∈ {0, 1})
2. (*User initialization*) Set initial index: 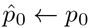
3. (*Recursive operation*) Set initial bit position: *k* = 0 **while** (*k* < *λ*) **do** 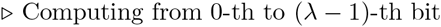

a. (*Query entry*) The user performs the following steps:

- Set an offset *o_k_*: **if** (*b^k^* (*c*) = 0) *o_k_* ← 0 **else** *o_k_* ← *N* + 1
- Calculate next index:

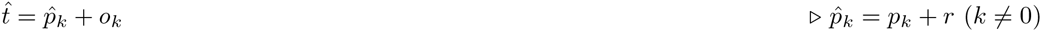
- Prepare next query:

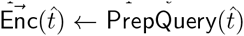
- Send 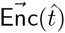 to the server.
b. (*Operate*) The server performs the following steps:

- Generate a random value *r*
- Set *r*′ ← 0 iff. *i* = 0
- Compute the next index:

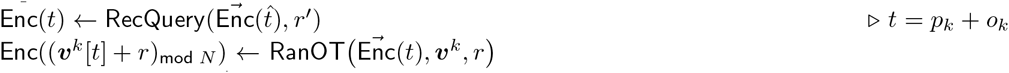
- Store a random value *r*′ ← *r*
- Send 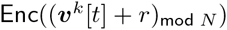 to the user
c. (*Receive randomized index*) The user obtains:

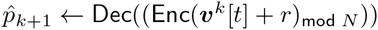

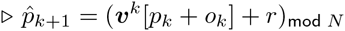 *k* ← *k* + 1 **end while**
4. The user sends the last query:

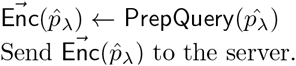
5. The server obtains encrypted *pλ*:

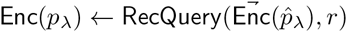

At the end of the protocol:

- The user holds: 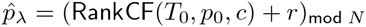
- The server holds: 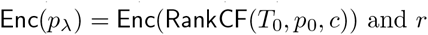

Those sub-modules are executed in the following order, and all steps are repeated until a condition predefined by the main algorithm calling `ROT` is satisfied.

**Step 1** The user conducts `PrepQuery` 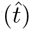 and sends 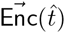 to the server.

**Step 2** The server conducts `RecQuery` 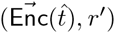 to obtain an encryption of a correct index 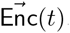.

**Step 3** The server conducts `RanOT` 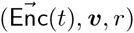 to compute an encrypted and randomized result 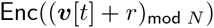.

**Step 4** The user obtains 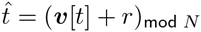 by `Dec`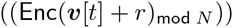 and goes back to **Step 1**.

`ROT` returns randomized results as long as the server uses *r* ≠ 0 in **Step 3**. For a full-text search, we use a trick for returning a proper flag to the user only when the longest prefix match is found, which is described in Section 3.4. See our previous work (Shimizu *et al.*, 2016) for more details about the `ROT` algorithm.

### 3.3 Secure Wavelet Matrix

Here, we describe the algorithm of sWM in detail. A pseudocode of sWM is written in Algorithm 1. The protocol starts with the initialization task, in which the server prepares sub-lookup tables ***υ***^0^,…, **υ**^*λ*−1^ (Step 1), and the user sets an initial index *p*_0_ (Step 2). *p* denotes *true* `RankCF`, which should be held by the user at the end of the protocol, and 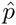 is the corresponding *randomized* `RankCF`, to which the server adds a random factor *r*. Analogously, we denote 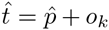 and *t* = *p* + *o_k_*, where *o_k_* is an offset for searching for ***υ***^k^. In the recursive search task (Step 3), the user and the server collaboratively compute 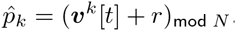. On the server side, the user’s input 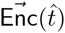 is *not* decrypted, and the result is computed by those ciphertexts. On the other hand, the user obtains a plaintext (***υ***[*t*] + *r*)_mod *N*_. However, this is randomized by *r*, and the user cannot see ***υ***[*t*] that includes the server’s private information. The random factor *r*is added by the ROT task in the third item of Step 3b. Therefore, the protocol is secure for both the user and the server sides. Though Algorithm 1 returns randomized results for all communication rounds, the user can obtain a true RankCF at the end of the protocol if the server uses *r* = 0 only at the final round of the protocol. In Step 4, the user sends an additional query to let the server hold an encrypted result `Enc`(*pλ*). This is an unnecessary task for computing `RankCF`(*T*_0_, *p*_0_, *c*), however, it is often convenient to hold an encrypted result `Enc`(*pλ*) by the server when the sWM is used as a building block of other search algorithm. We will introduce such a search algorithm in Section 3.4, and will show how the encrypted result held by the server is used in the algorithm.

### 3.4 Secure FM-Index

In this study, we apply the sWM to the problem of full-text search by using the FM-Index algorithm, in which a sorted string called Burrows Wheeler Transform (BWT) is created for indexing a database string (Ferragina and Manzini, 2000). As we mentioned in Section 2.2, a substring match is reported as an interval [*f, g*) on the BWT, and there is a recursion relationship described in Equation (1). (For the case of FM-Index, *%* in Equation (1) is BWT.) The main idea of our algorithm is to use sWM to compute `RankCF`. It repeats sWM until the longest match is found (e.g., *g* = *f*). Algorithm 2 is the detailed algorithm of the secure FM-Index (sFMI). Note that some of the steps in Algorithm 1 are slightly modified to fit it to the FM-Index algorithm. The key part of the sFMI is in Step 3b, in which the server computes an encrypted flag by using the function isLongest. This flag becomes an encrypted 0 only when the match is longest, and becomes an encryption of a random value otherwise. Therefore, only the user knows whether or not each substring match is the longest by checking the flag in Step 3c. The algorithm is scalable for different search options if the function isLongest is replaced by another function holding a different end condition. For example, it enables searching for a longest substring match whose occurrence is at least *E*, just by computing encrypted flags 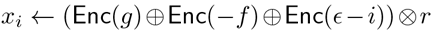 for *i* = 0,…, ϵ−1 and checking whether or not it includes an encrypted 0. (The algorithm is detailed in Section S1 in the supplementary material.) Since the algorithm repeats sWM **ℓ** times for the query of length **ℓ**, total time complexity becomes 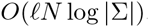. We will discuss the complexity in more detail in Section 3.5.

The original FM-Index algorithm provides a backward search where a search direction starts from the tail to the head of a query. This can be easily converted into a forward search by querying in the reverse direction. For simplicity, we described the algorithm in such a way that the query is searched in a forward direction.

### 3.5 Complexity

In this section, we compare the theoretical complexity of sFMI and PBWT-sec. The computational obstacle of both algorithms is a ROT task. Given a lookup-table of length *N*, the time complexity of the `ROT` is *O_N_* and the communication complexity is *O*(*√N*). It might not be intuitive that the communication complexity is not linear to the size of the lookup-table. However, there is an efficient algorithm in which each item of a lookup table is described in two-dimensional form that enables the length of a query to be reduced to *√N*. See our previous work (Shimizu *et al.*, 2016) for more detailed information.

The sWM used in sFMI repeats `ROT` for all sub-lookup tables, and thus the time and communication complexities become *O*(*N* log |Σ|) and *O*(*√N* log |Σ|), respectively. On the other hand, PBWT-sec performs a single ROT task for a lookup-table of length *N*|Σ|. Therefore, its time complexity is *O*(*N*|Σ|) and the communication complexity is *O*(*√N*|Σ|). Apparently, time complexity is an order of magnitude better for s`FMI`,while the communication complexity is comparable as long as the Σ is not very large. Since the communication complexity is already sub-linear to the size of *N*, the improvement in time complexity is much more important for practical problems.

Table 1 summarizes time and communication complexities of both sFMI and PBWT-sec. Note that both time and communication complexities are multiplied by the query length **ℓ**. In addition to those two algorithms using a clever data structure, for comparison, we also describe complexity of a Na¨ıve method, in which all occurrences of substrings appearing in a database text are stored in a lookup table and the user uses ROT to search for a substring match.

**Table 1:**
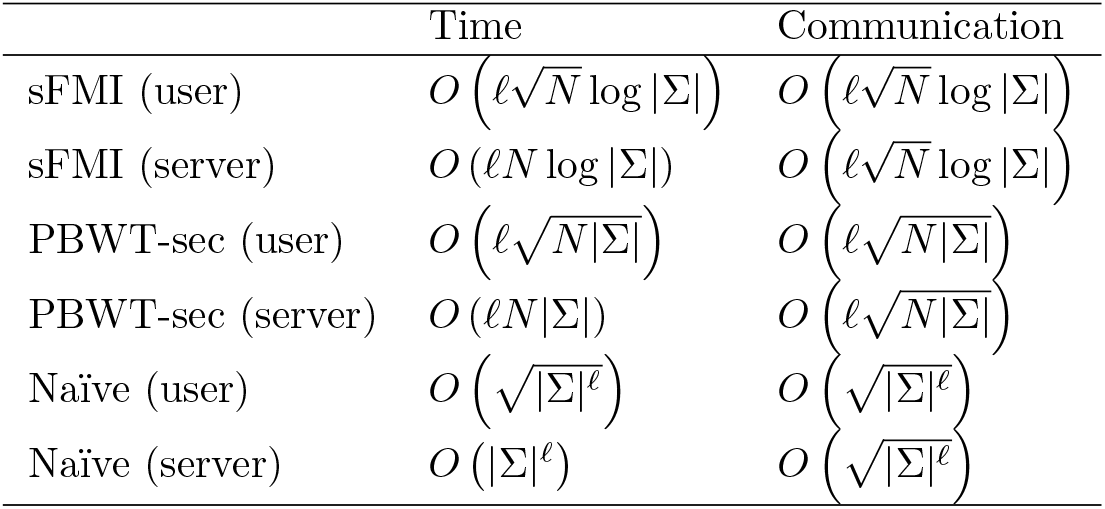
Summary of time and communication complexities of sFMI (proposed method), PBWT-sec (previous method), and Na¨ıve approach.

## 4 Experiments

To evaluate the efficiency of the proposed method (sFMI), we performed experiments on both a simulated dataset and two different real datasets. For the simulated dataset, we simply generated random strings each of which has a length of 10,000 with |Σ| = 4,8,16,…,1024. For the real datasets, we used all the protein sequences included in Ribosomal S4Pg Family of Pfam (Finn *et al.*, 2016) and all clinical study titles (879 titles, in Japanese, 53,560 characters in total excluding punctuation) stored in JAPIC Clinical Trials Information (JAPIC, 2008). For each real dataset, all the sequences are concatenated into a long single sequence with a delimiter symbol. The Japanese text is usually written in a combination of a Japanese alphabet (Hiragana, |Σ|= 83 including sound marks) and Chinese ideographs (|Σ| > 10,000). However, Chinese ideographs can be spelled out into Hiragana, though this is generally very unnatural. Therefore, we prepared two datasets for the clinical text: one is the converted text written in Hiragana (Clinical DB1), and the other is the original text (Clinical DB2). Those texts also include words written in Arabic numerals (|Σ|= 10) and Roman (case-sensitive) and Greek (case-insensitive) alphabets (|Σ|= 26, 50 respectively), because numbers and technical terms including Roman/Greek letters are usually written in their original form. Clinical DB1 consists of the alphabet of |Σ|= 170, and Clinical DB2 consists of the alphabet of |Σ|= 21,207 including the delimiter symbol. (Since one Chinese ideograph is usually spelled out by more than one Hiragana character, Clinical DB1 and DB2 have exactly the same meaning but different lengths.) See Section S2 in the supplementary material for more details about the character sets.

We implemented the proposed algorithm in C++ based on an open source C++ library of *elliptic curve ElGamal encryption* (Mitsunari, 2016). We used the same implementation of ROT for all the three methods, which is provided by another C++ library (Shimizu, 2016). For the security parameters, we used a standard configuration called *secp192k1* (SECG curve over a 192-bit prime field) in accordance with the recommendation by The Standards for Efficient Cryptography Group. We used a laptop computer equipped with Corei7 3.00GHz (2 physical cores) for the user, and a standard desktop PC equipped with Xeon 3.40GHz (12 physical cores) for the server for all of the experiments. For the simulated dataset, the program was run with 2 threads for the user and 8 threads for the server. For the read datasets, it was run with 2 or 4 threads for the user, and 8 or 16 threads for the server.

Figure 5 shows run time and communication size for the simulated dataset when query length is 10. As shown in panel (a), an observed run time of sFMI is order of magnitude faster than PBWT-sec, which is concordant with the theoretical complexity. Communication size of the proposed method is also better than that of PBWT-sec when the |Σ| is sufficiently large.

**Figure 5:**
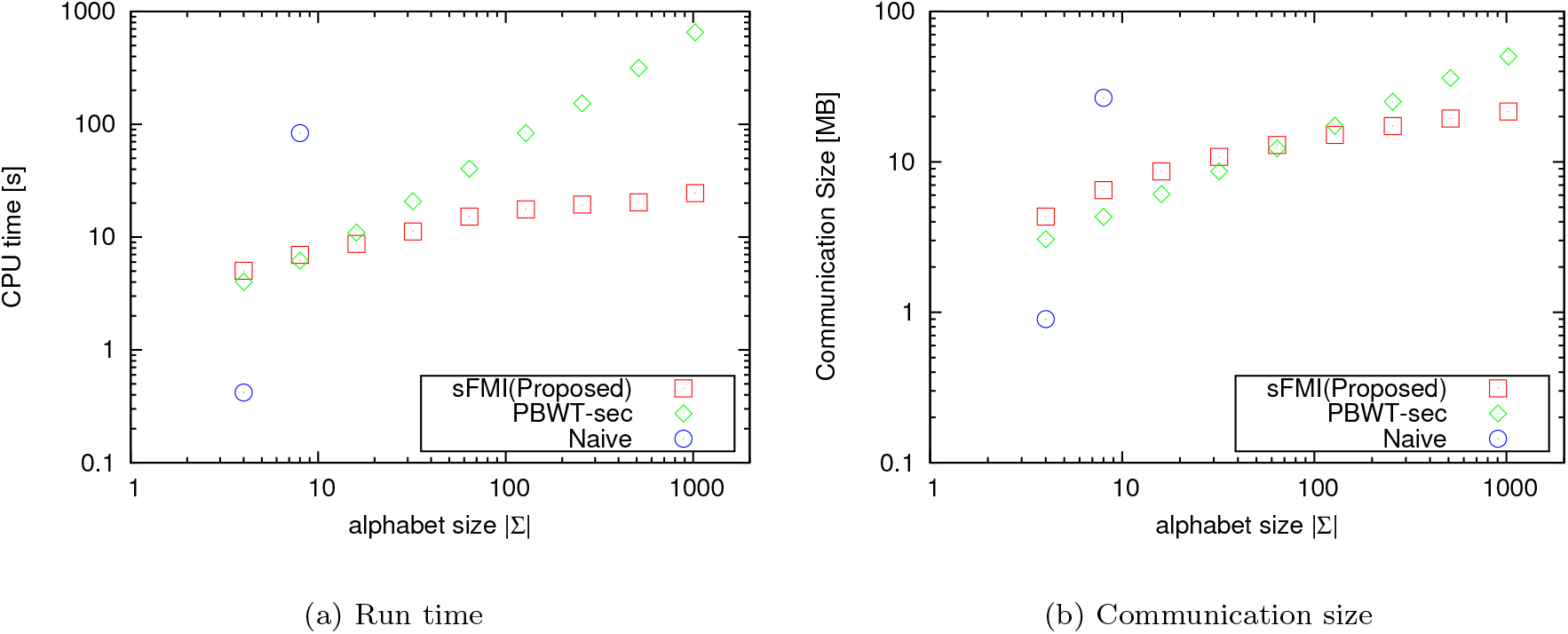
(a) Run time and (b) Communication size of sFMI, PBWT-sec, and Naiv¨e method for random string of 10,000 with alphabet size |Σ|= 4,8,16,…, 1024, when query length is 10. Server used eight threads, and user used two threads. Run time includes communication overhead.

Table 2 shows run time for searching on real datasets. For the search on Clinical DB1 described by the Hiragana alphabet, server-side run time of the proposed method was 10 times faster than that of the previous method. For the search on Clinical DB2, server-side run time of the proposed method is only 103.74 although |Σ| is huge, while the search by the previous approach did not finish within 24 hours. The results show that our method is practical even for real datasets.

**Table 2:** Run time of sFMI (proposed method) and PBWT-sec (previous method) for searching on protein sequence database (protein DB) and clinical study results database described in reduced set of Hiragana alphabet (Clinical DB1) and that in original form (Clinical DB2).

|  | Protein DB |  | Clinical DB1 |  | Clinical DB2 |  |
| --- | --- | --- | --- | --- | --- | --- |
| Server run time (s): |  |  |  |  |  |  |
| Num. of threads | 8 | 16 | 8 | 16 | 8 | 16 |
| sFMI | 8.28 | 6.66 | 75.93 | 59.72 | 103.74 | 80.00 |
| PBWT-sec | 12.54 | 9.94 | 856.78 | 554.40 | - | - |
| User run time (s): |  |  |  |  |  |  |
| Num. of threads | 2 | 4 | 2 | 4 | 2 | 4 |
| sFMI | 2.56 | 2.50 | 9.32 | 9.20 | 15.28 | 14.65 |
| PBWT-sec | 1.27 | 1.21 | 9.65 | 9.55 | - | - |
| DB Length | 9,826 |  | 77,712 |  | 53,560 |  |
| Alphabet Size | 21 |  | 170 |  | 21,207 |  |
| Query length | 10 |  | 10 |  | 10 |  |

## 5 Conclusion

We have developed an efficient algorithm for a secure string search on the basis of a novel technique combining wavelet matrix and homomorphic encryption. It can search any type of string while still protecting privacy as strongly as the previous approach. We implemented the proposed method and tested it on both a simulated dataset and real datasets. The results show that the proposed method is an order of magnitude more efficient than the previous approach in terms of alphabet size and that its computational cost is acceptable for practical use. As the proposed method potentially scales to various types of data such as two-dimensional data and tree data, it is expected to be used for an even wider range of life science data and contribute to secure data sharing.

## Algorithm 2

Detailed description of secure FM-Index.

**protocol** `sWM` 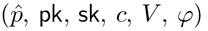

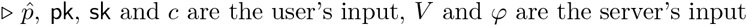

Perform Algorithm 1 with the following modifications

- Use `pk` and `sk` for a public key and a private key
- Omit Step1 and use input *V* as sub-lookup tables
- Initialize 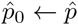 in Step2
- Initialize 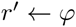 in the second item of Step 3b

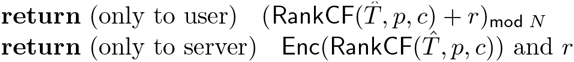

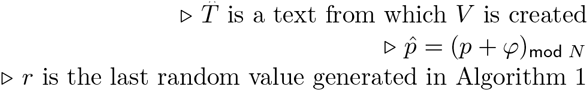

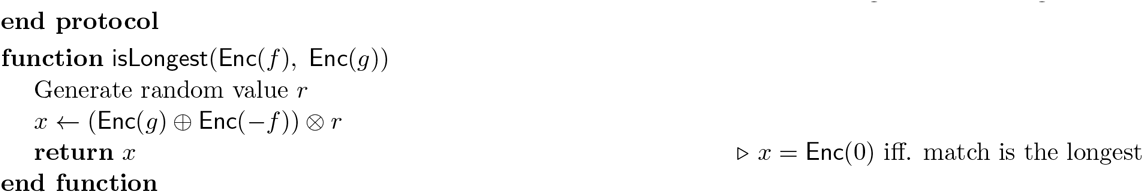
- Public input: Problem size *N*; alphabet Σ
- Private input of user: A query sequence ***q*** of length **ℓ**
- Private input of server: A database text *T*

0. (*Key setup of cryptosystem*) The user generates key pair (`pk, sk`) by using key generation algorithm KeyGen for additive-homomorphic cryptosystem and sends public key pk to server

1. (*Server initialization*)

- The server creates BWT of *T* and stores it as 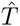
- The server creates a set of sub-lookup tables for 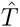:
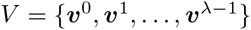, by using the same process described in Step 1 of Algorithm 1
2. (*User initialization*) Set initial interval 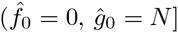
3. (*Recursive search*) Initialize an index: 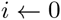 Initialize random factors: 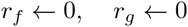

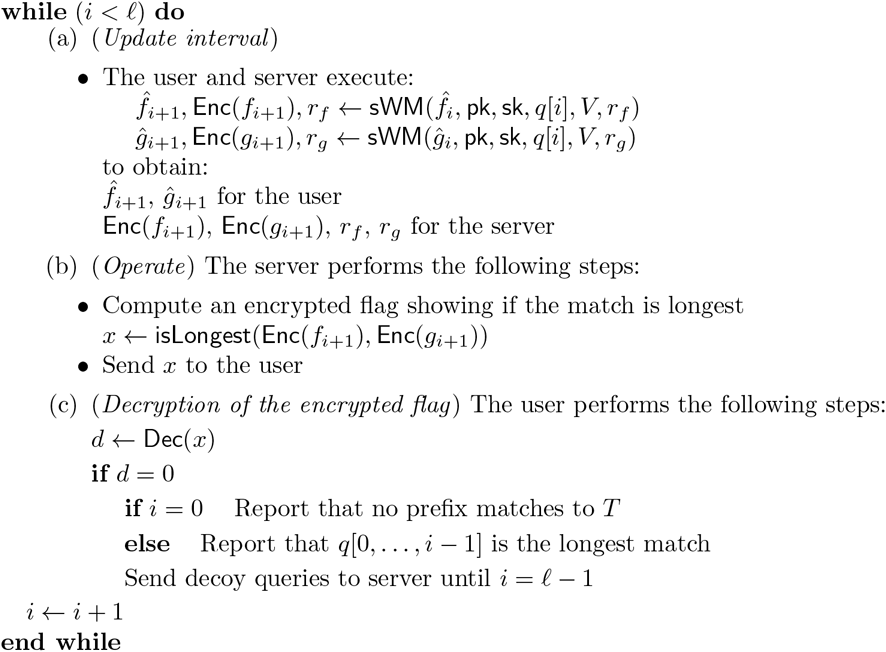

The user reports that 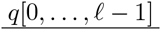 is the longest match, if 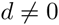 for *i* = 0,…, *ℓ* − 1.

## S1 Additional search option

We referred to different search options of the secure FM-Index in Section 3.4. Here, we describe in detail one of those search options that enables searching for the longest substring match whose occurrence is at least *ϵ*. A server can avoid leaking information about rare substrings in its database to a user by using this search option. Recall that in FM-Index, *k*-th reported intervals [*f_k_, g_k_*) implies k-prefix of a query matches to the database with *g_k_ −f_k_* positions. Therefore, checking whether occurrence of substring is less than *ϵ* is equivalent to *g−f − ϵ* < 0.The key idea of implementation is the design of flags: each flag indicates whether *g − f − i*(*i* = 0,1,…, *ϵ* − 1) is 0 or not. *i*-th flag is 0 iff. k-prefix of a query matches to the database with *i* positions. Therefore, only one flag will be 0 iff. *g − f − ϵ* < 0. In practice, all flags are randomized and encrypted.

The outline is as follows:

> **Server side procedure**
>
> The server prepares encrypted flags ***x*** as follows:
>
>
> 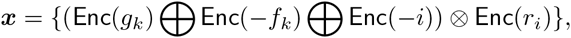
>
>
> where *i* = 0,1,…,*ϵ* − 1, each *r_i_* is a random value different from the other *r_j_*. The server shuffles and sends ***x*** to the user.
>
> **User side procedure**
>
> The user then decrypts ***x*** and checks whether one of the flags is 0 or not (only one flag will be 0 at most). If one of the flags is equal to `Enc(0)`, the user knows the occurrence of *k*-prefix substring match is less than *ϵ*. Note that the user cannot know the exact occurrence of a substring match.

To implement this search option, we replace `isLongest` with another function `isELongest` corresponding to the server side procedure above. Also, we need to slightly modify the user side procedure for checking the end condition. A detailed algorithm of modified secure FM-Index is presented in Algorithm S3. isELongest function and Step 3b are mainly modified parts.

### Algorithm S3

Detailed description of secure FM-Index.

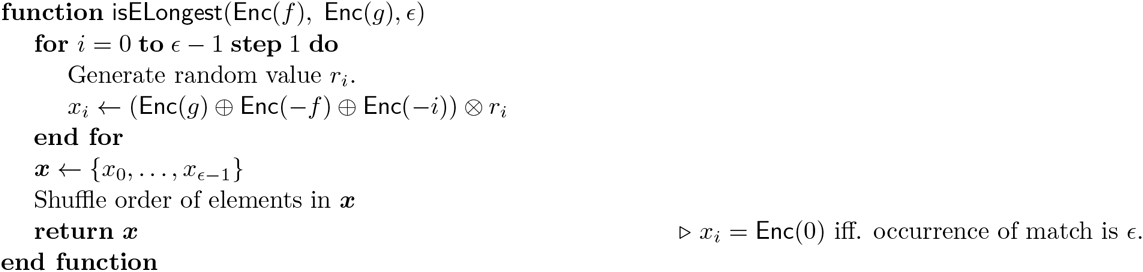

- Public input: Problem size *N*; alphabet Σ
- Private input of user: A query sequence ***q*** of length **ℓ**
- Private input of server: A database text *T*

0. (*Key setup of cryptosystem*) User generates key pair (`pk, sk`) by key generation algorithm KeyGen for additive-homomorphic cryptosystem and sends public key `pk` to server.

1. (*Server initialization*)

- Server creates BWT of *T* and stored it as 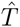.
- Server creates a set of sub-lookup tables for 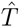: 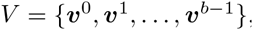, by the same process described in Step 1 of Algorithm 1
2. (*User initialization*) Set initial interval 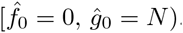.
3. (*Recursive search*) Initialize an index: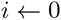 Initialize random factors: 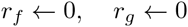

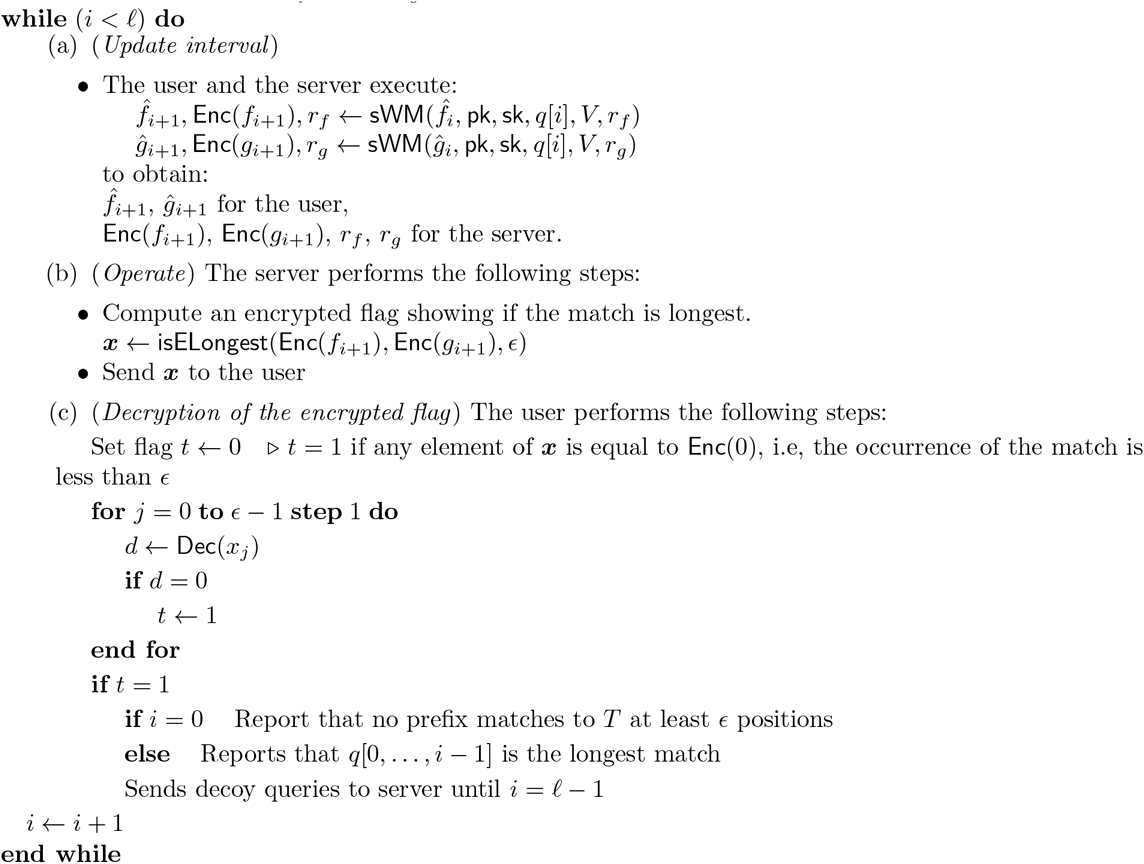

The user reports that 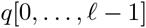 is the longest match, if 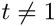 for *i* = 0*,…, *ℓ* −* 1.

## S2 Characters used in the experiments

Table S1 shows a set of characters and corresponding code points of Unicode that were used in the experiments. We used a CJK unified ideographs table which is included in Unicode version 8.0, because it contains most of the Chinese ideographs that are commonly used in Japan.

**Table S1:** Unicode code points of characters included in Clinical DB1 and DB2

|  | Unicode code point | DB name |
| --- | --- | --- |
| Arabic numerals | 0x0030 - 0x0039 | Clinical DB1, 2 |
| Roman alphabet (lower case) | 0x0041 - 0x005A | Clinical DB1, 2 |
| Greek alphabet (upper case) | 0x0391 - 0x03A9 | Clinical DB1, 2 |
| Greek alphabet (lower case) | 0x03B1 - 0x03C9 | Clinical DB1, 2 |
| Hiragana | 0x3041 - 0x3093 | Clinical DB1, 2 |
| Symbols for long vowel sound | 0x30FC | Clinical DB2 |
| Katakana | 0x30A1 - 0x30F6 | Clinical DB2 |
| Chinese ideograph (CJK unified ideographs table) | 0x4E00 - 0x9FD5 | Clinical DB2 |

